# Generating an Artificial Nest Building Pufferfish in a Cellular Automaton Through Behavior Decomposition

**DOI:** 10.1101/478750

**Authors:** Thomas E. Portegys

## Abstract

A species of pufferfish builds fascinating circular nests on the sea floor to attract mates. This project simulates the nest building behavior in a cellular automaton using the Morphognosis model. The model features hierarchical spatial and temporal contexts that output motor responses from sensory inputs. By considering the biological neural network of the pufferfish as a black box, decomposing only its external behavior, an artificial counterpart can be generated. In this way a complex biological system producing a behavior can be filtered into a system containing only functions that are essential to reproduce the behavior. The derived system not only has intrinsic value as an artificial entity but also might help to ascertain how the biological system produces the behavior.

## Introduction

A species of pufferfish creates an astounding circular nest structure on the sea floor near Japan, as shown in Figure 1. The nest is built by the male for the purpose of attracting a female for mating. The nest contains a central smooth area surrounded by radial furrows and is about 2 meters in diameter. The 12 centimeter long pufferfish sculpts the furrows by sweeping sediments with its fins as its travels back and forth lengthwise along each furrow. The female lays her eggs in the fine sediments in the central area which are then fertilized by the male (Kawase et al., 2013).

**Figure 1.**
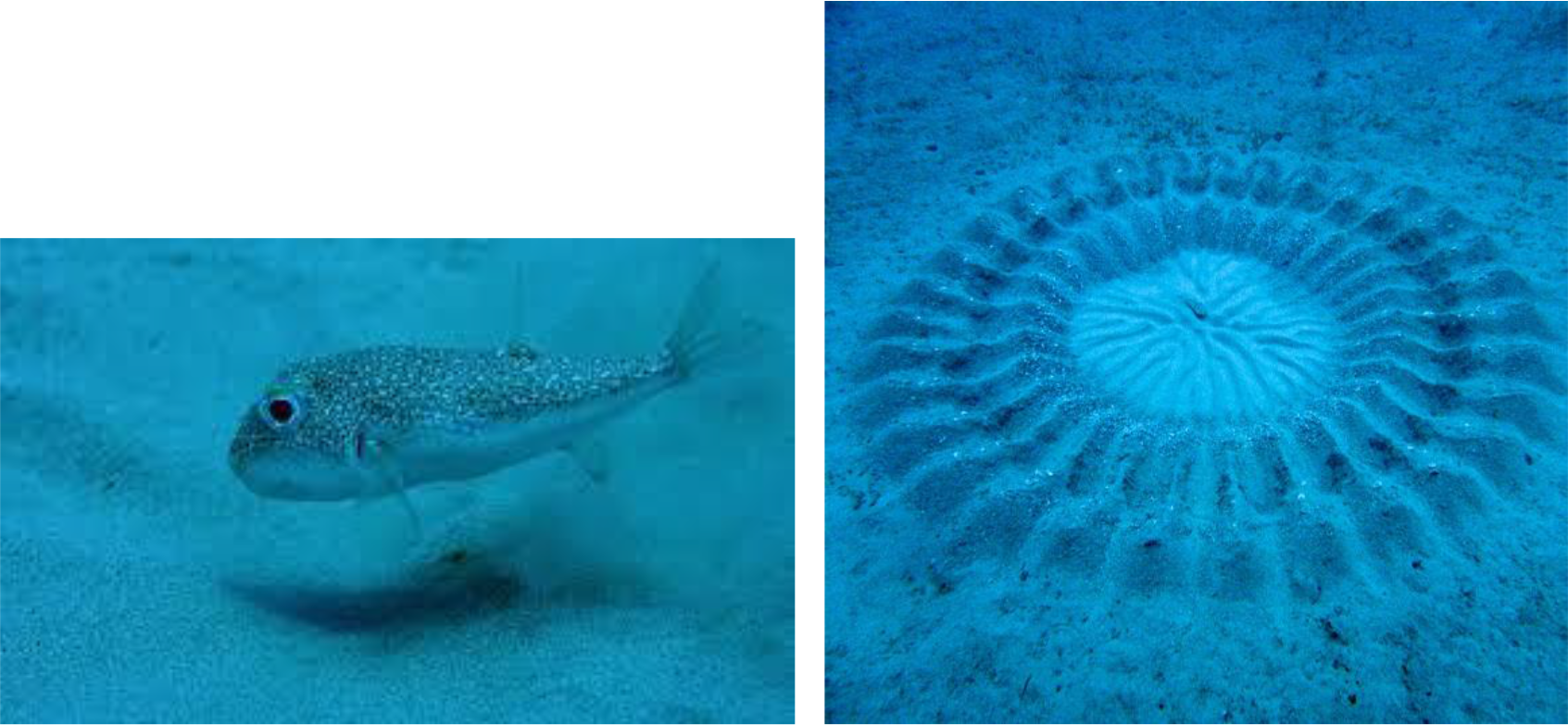
Pufferfish (left), and nest (right).

How does a pufferfish, whose brain contains approximately 10 million neurons, instinctively construct such a complex structure? Nature is rife with such interesting animal exploits. Termites build and maintain complex nests by coordinating the efforts of a massive number of individuals. Spiders build complex webs. Birds construct elaborate nests. Honey bees coordinate foraging through a body language signaling system. These organisms are all related to humans to some degree. Considering nature’s propensity for extending and repurposing capabilities, unlocking the secrets of animal intelligence is a worthy step toward understanding the underpinnings of human intelligence.

Evolution is a trial-and-error tinkerer that makes extremely complex life forms. Even the well-known tiny nematode worm, C. elegans, with only approximately 1000 cells, 302 of which are neurons, remains a daunting subject for researchers. On an even smaller scale, geneticists are beginning to entertain the notion that most traits are the result of contributions from a great deal of the genome (Boyle et al., 2017).

One proposal to extract essential information from nature’s witch’s brew of complexity is the “radical reimplementation” approach (Lehman and Stanley, 2014). The idea is to discover general principles by intentionally diverging from nature in features that might be incidental or artefactual. If the core behavior is preserved after diverging, then the reimplemented features can be excluded from general principles. As an example, consider the case for discovering principles of flight from examination of bird wings. In the biological implementation, flapping wings are universal; yet in the artificial realm, fixed wings are the norm. It is the aerodynamics of wings that is common to both.

In simpler creatures reverse engineering is sometimes feasible. For example, the navigation skills of honey bees are of value to drone technology. Fortunately, it appears that the modular nature of the honey bee brain can be leveraged to replicate this skill (Nott, 2018).

Another approach is to consider a biological system as a black box and decompose its behavior into components that model the behavior in an artificial implementation. The expectation is that the artificial system will be less complex than the biological one, capturing only what is essential to recreate the behavior, and possibly more amenable to analysis. For artificial intelligence, which is more concerned with function over form, this can be an end in itself. Yet even for biologists, the artificial system may offer hints as to how the biological system performs.

An example of this approach addresses a need to distinguish mutant strains of C. elegans nematode worms based on movement variations (Li et al., 2017). An artificial neural network (ANN) was trained to closely emulate recorded worm trajectories. Although the network was not directly trained to perform classification, neuron responses within the network were found to perform well as mutant strain classifiers.

This project simulates pufferfish nest building behavior in a cellular automaton using the Morphognosis model. The model features hierarchical spatial and temporal contexts that output motor responses from sensory inputs. Through a black box decomposition of nest building behavior into space-time rules, an artificial pufferfish is embodied an ANN. This process is depicted in Figure 2.

**Figure 2.**
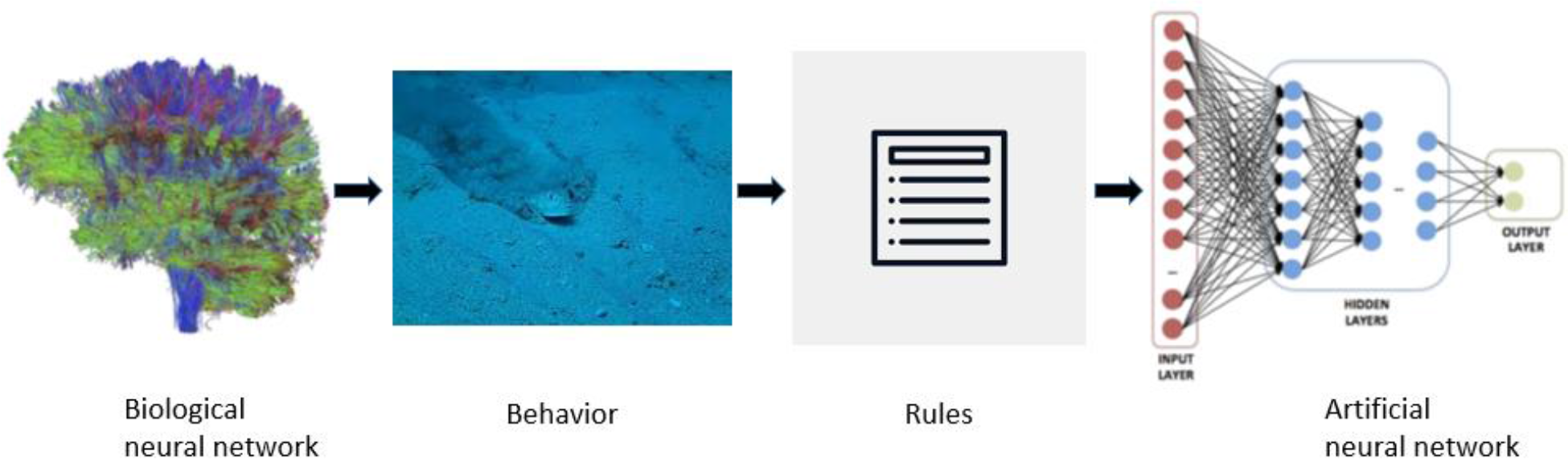
Behavior decomposition and artificial neural network reconstitution.

Pufferfish nest building has been accurately simulated by Mizuuchi et al. (2018) in a manner that bears close resemblance to the actual nest. Here, in contrast, nest building was selected as a complex task to be learned by Morphognosis rather than an end in itself, and thus the simulation is more abstracted.

## Description

### Morphognosis Overview

The brain, a complex structure resulting from millions of years of evolution, can be viewed as a solution to problems posed by an environment existing in space and time. Internal spatial and temporal representations allow an organism to navigate and manipulate the environment. Neurological research has confirmed that animals have specific structures that process spatial and temporal information (Hainmüller and Bartos, 2018; Lieff, 2015; Vorhees and Williams, 2014).

These concepts are embodied in a model called *Morphognosis* (*morpho* = shape and *gnosis* = knowledge). Introduced with several prototype tasks (Portegys, 2017), Morphognosis has also modeled the locomotion and foraging of the C. elegans nematode worm (Portegys, 2018).

#### Morphognostics

The basic structure of Morphognosis is a pyramid of event recordings called a *morphognostic*, shown in Figure 3. At the apex of the pyramid are the most recent and nearby events. Receding from the apex are less recent and possibly more distant events. A morphognostic can thus be viewed as a structure of progressively larger nested chunks of space-time knowledge that form a hierarchy of contexts. A set of morphognostics forms long-term memories that are learned by exposure to the environment. Scaling is accomplished by aggregating event information. This means that more recent and nearby events are recorded in greater precision than events more distant in space and time.

**Figure 3.**
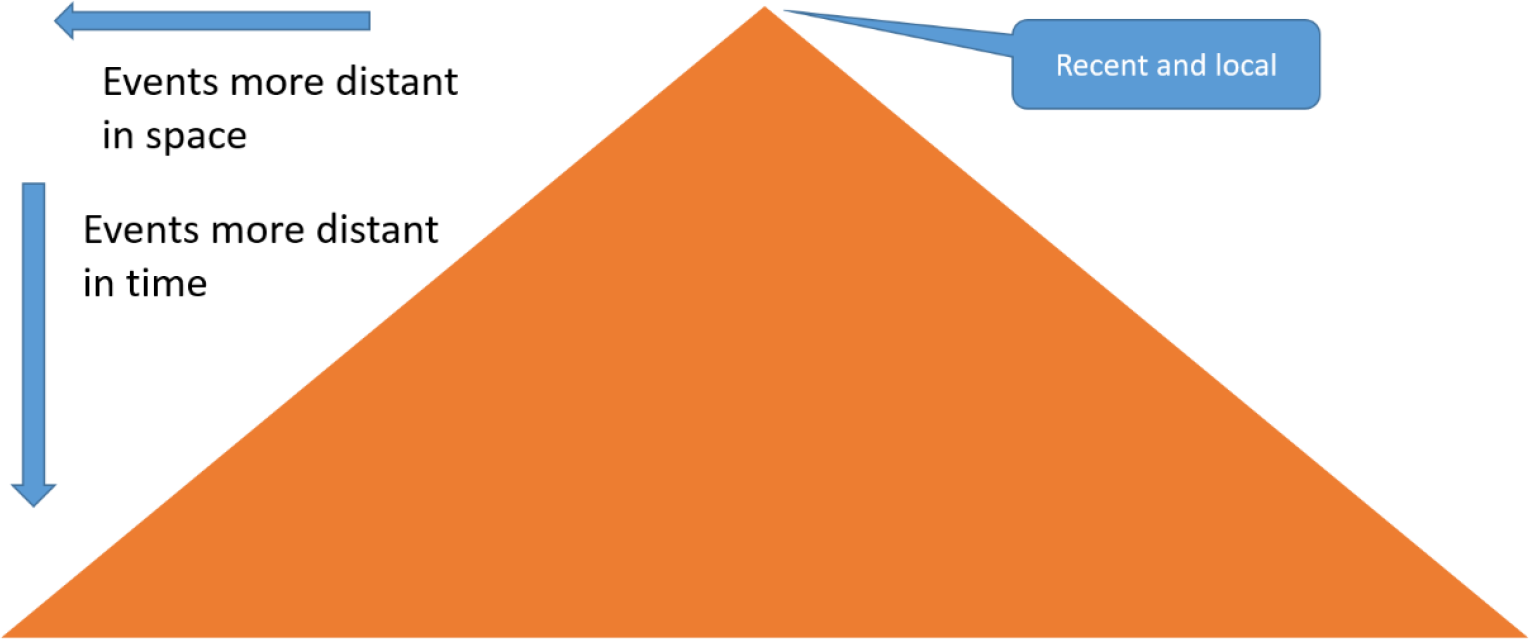
Morphognostic event pyramid.

Figure 4 is an example of a morphognostic implemented in a cellular automaton as a nested set of 3×3 neighborhoods and aggregated histograms of cell state densities.

**Figure 4.**
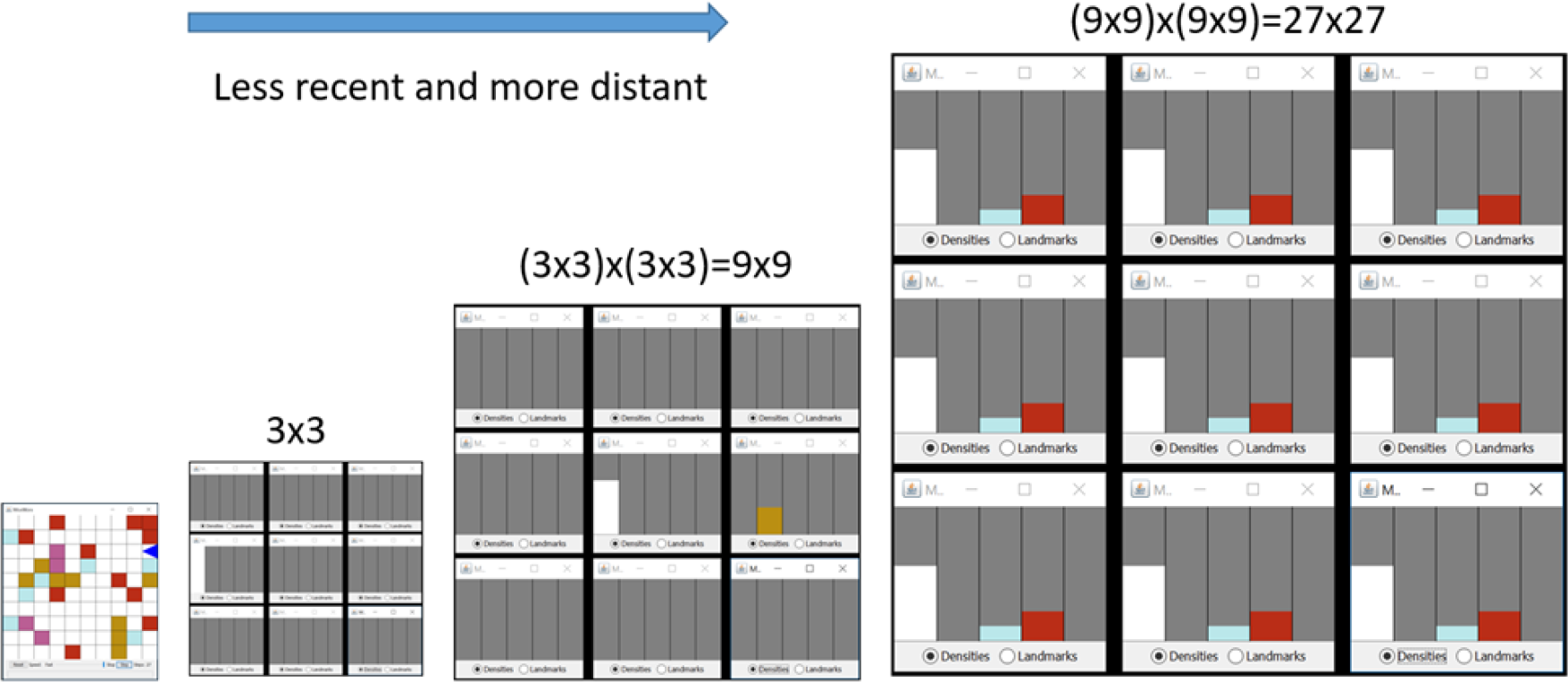
Cellular automaton implementation of Morphognosis.

The following are formal definitions of the spatial and temporal morphognostic neighborhoods.

##### Morphognostic Spatial Neighborhoods

A cell defines an elementary neighborhood:

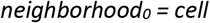

A non-elementary neighborhood consists of an *N×N* set of *sectors* surrounding a lower level neighborhood:

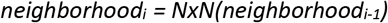

where *N* is an odd positive number.

The value of a sector is a vector representing a histogram of the cell type densities contained within it:

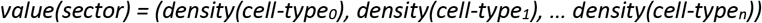

The number of cells contributing to the density histogram of a sector of *neighborhood*_*i*_ = *N*^*i-1*^*×N*^*i-1*^

##### Morphognostic Temporal Neighborhoods

A neighborhood contains events that occur between time *epoch* and *epoch* + *duration*:

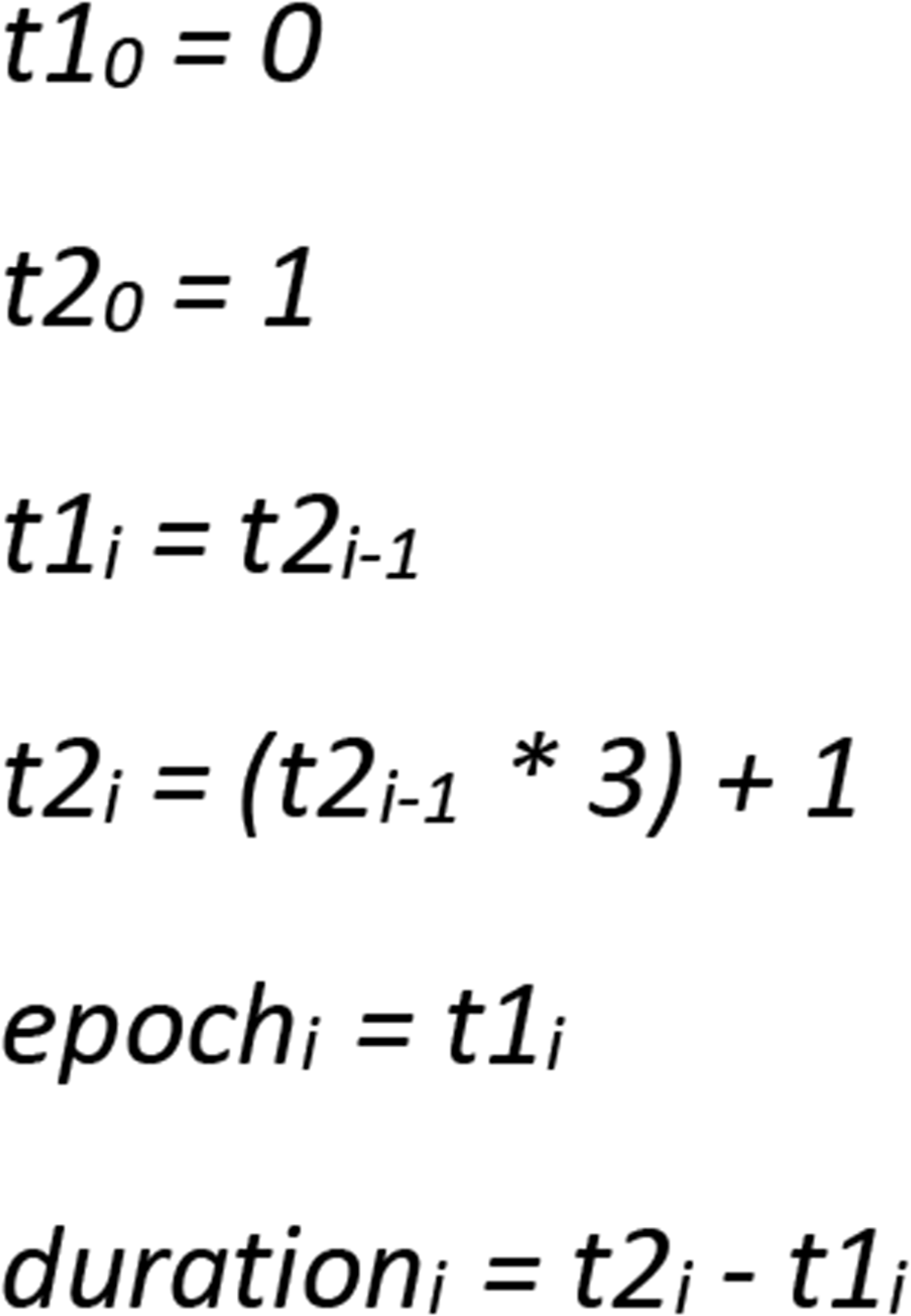

#### Metamorphs

In order to navigate and manipulate the environment, it is necessary for an agent to be able to respond to the environment. A *metamorph* embodies a morphognostic→response rule. A set of metamorphs can be learned from a manual or programmed sequence of responses within a world.

Metamorph “execution” consists generating a morphognostic for the current environmental state and then finding the closest morphognostic contained in the learned metamorph set, where:

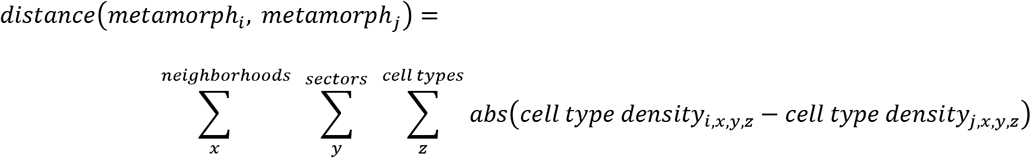

#### Artificial Neural Network Implementation

In a complex environment, generating a large number of metamorphs may be prohibitive in terms of storage and search processing. Alternatively, metamorphs can be used to train an ANN, as shown in Figure 5, to learn responses associated with morphognostic inputs. During operation, a current morphognostic can be input to the ANN to produce a learned response. The ANN also has these advantages:

- Faster.
- More compact.
- Noise tolerance and generalization through weighting of information by importance.

**Figure 5.**
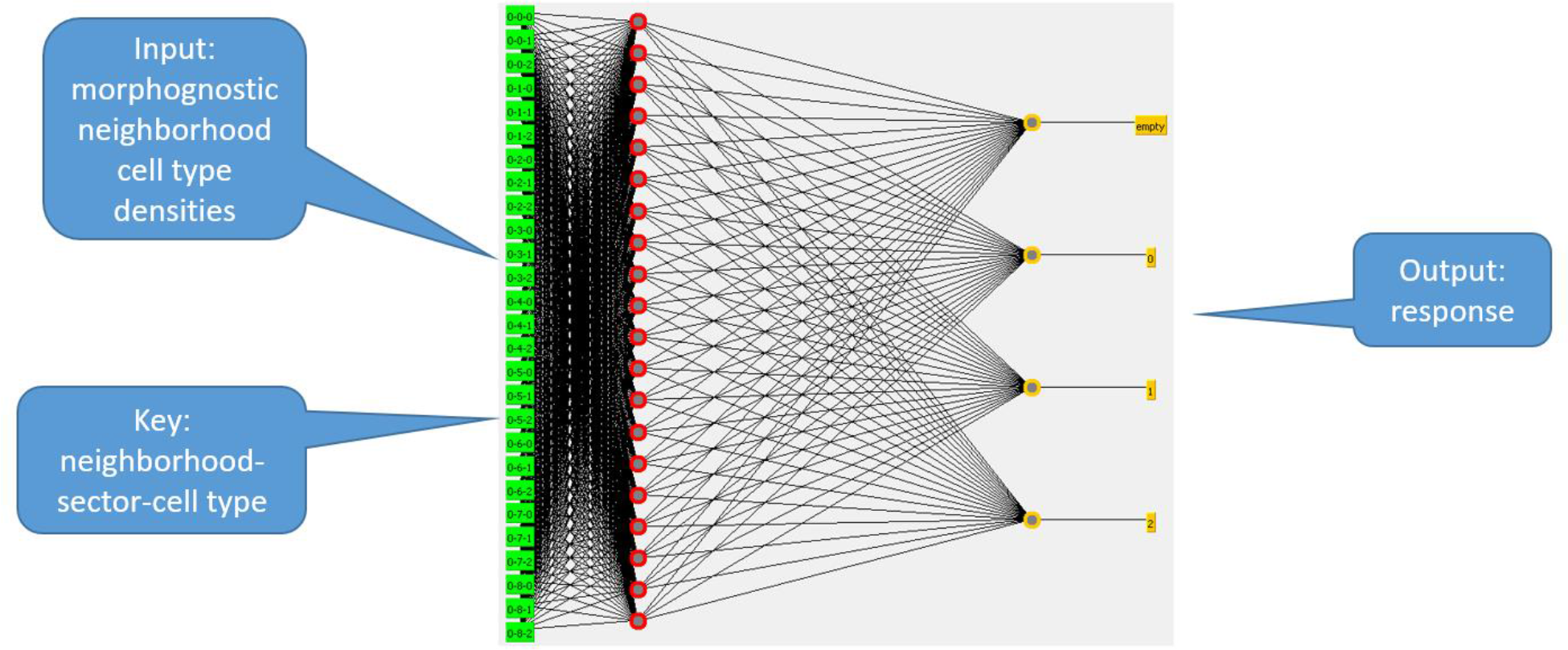
Metamorph artificial neural network.

## Pufferfish Nest Building

The pufferfish constructs its nest by first smoothing out a central area through the action of its fins. It then typically creates 28 radial furrows or spokes emanating from the central area by moving back and forth along them. A furrow has a set of dips and rises, produced by the pufferfish lingering in specific portions of the furrow while its fins disperse sediments outward. The result is shown in Figure 1.

A cellular automaton (CA) (Toffoli and Margolus, 1987) is used as a platform for the nest building. A CA is a grid of cells that contain state values. CAs are ideal for investigating systems that have parallel local production rules that output actions as a function of local cell states. For the pufferfish project, cells contain a surface elevation value. The surface is given a random initial set of elevations to simulate sea floor irregularities.

The pufferfish occupies a single cell and has an orientation of north, south, east and west. It can sense the elevations of three cells in front of it: left, center, and right. It can also sense its previous response. A set of sensor values constitutes an event. These are its responses: wait, forward, turn left, turn right, smooth sensed surface cells, raise and lower surface at current location.

The metamorph production rules are generated by recording the activity of an “autopilot” pufferfish. The autopilot performs a simple reenactment of the natural pufferfish’s nest building activity: smooth the central area, then sequentially create the radial furrows by raising and lowering the surface appropriately. The software allows a number of parameterized variations, including the size of the central area and the number and length of the furrows. As the autopilot proceeds, morphognostic/response pairs are encoded into metamorphs. Once the nest is built, the environment is reset and the pufferfish runs entirely on its metamorph productions.

For successful nest building, the Morphognosis parameters are set to:

`NUM_NEIGHBORHOODS=4`
`NEIGHBORHOOD_INITIAL_DIMENSION=3`
`NEIGHBORHOOD_DIMENSION_STRIDE=0`
`NEIGHBORHOOD_DIMENSION_MULTIPLIER=3`
`EPOCH_INTERVAL_STRIDE=1`
`EPOCH_INTERVAL_MULTIPLIER=3`

This means that spatially there are 4 nested neighborhoods that span cell extents of 3×3, 9×9, 27×27, and 81×81. Temporally, the neighborhoods contain event information of 0/1, 1/3, 4/9, and 13/27, where each pair is an epoch/duration. So for example, the second neighborhood contain event information within a 9×9 set of cells that occurred between times 1 and 3.

A neural network can be trained from the metamorphs by flattening them into a dataset of sensory features association with a response. The neural network compresses the production rules for faster execution. It also can be used to generalize: a number of cell surface elevation variations can be trained together that allows the nest to be built regardless of initial surface elevation configurations.

A manual control of the pufferfish is also provided. This allow a user to acquire a set of metamorphs that capture an arbitrary sequence of events.

As shown in Figure 6, a dashboard allows a user to observe the sensors and response of the simulated pufferfish, select the driver: autopilot, metamorph ruleset, or manual, and display the current morphognostic.

**Figure 6.**
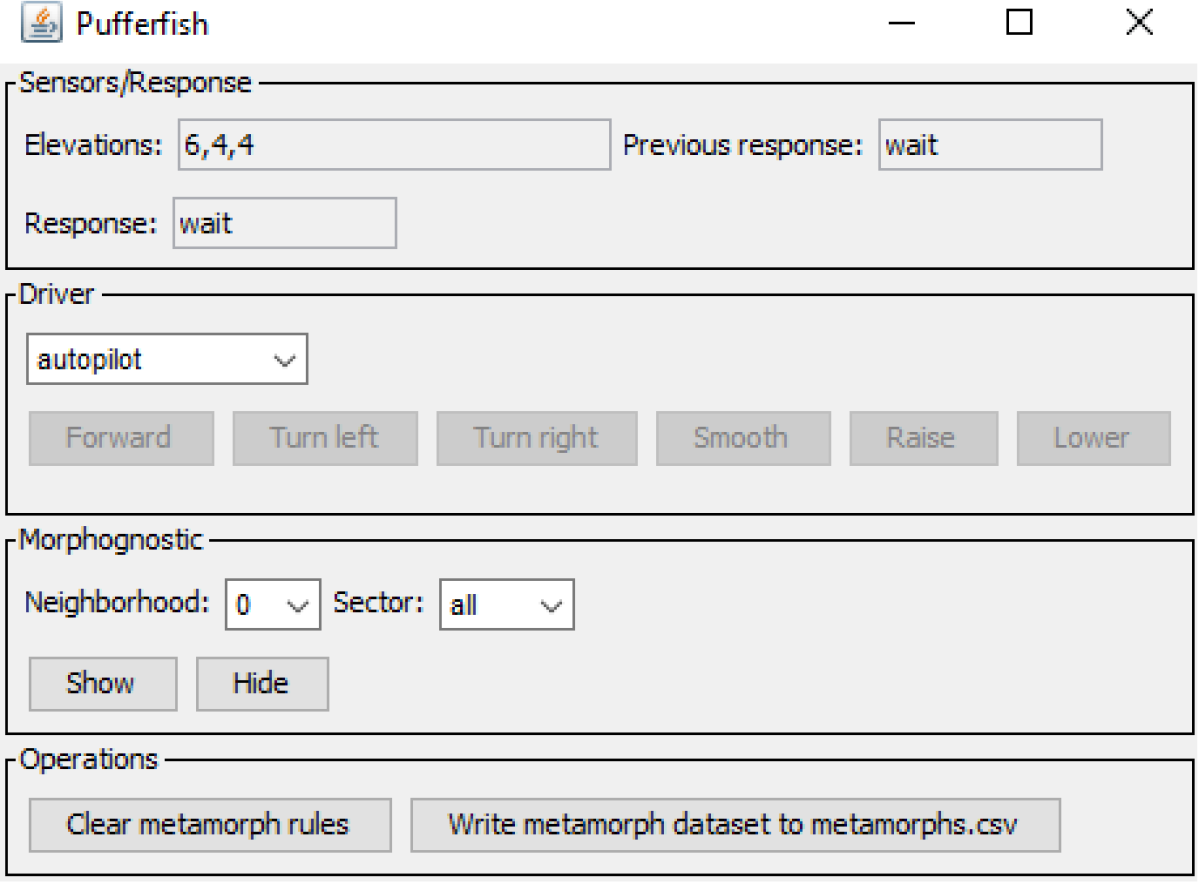
Pufferfish dashboard.

The Java code is available at: https://github.com/morphognosis/Pufferfish. The software is prebuilt and requires only unpacking and a single click to run.

## Results

Using the procedure described above, the pufferfish is capable of building nests of various sizes and shapes. Figure 7 shows such a nest on a 21×21 grid. There are 28 radial furrows which overlap due to the cellular grid, and each furrow replicates the down-up-down-up elevation sculpting of the sediments that exists in the actual nest. To build the nest, 903 metamorph rules were acquired.

**Figure 7.**
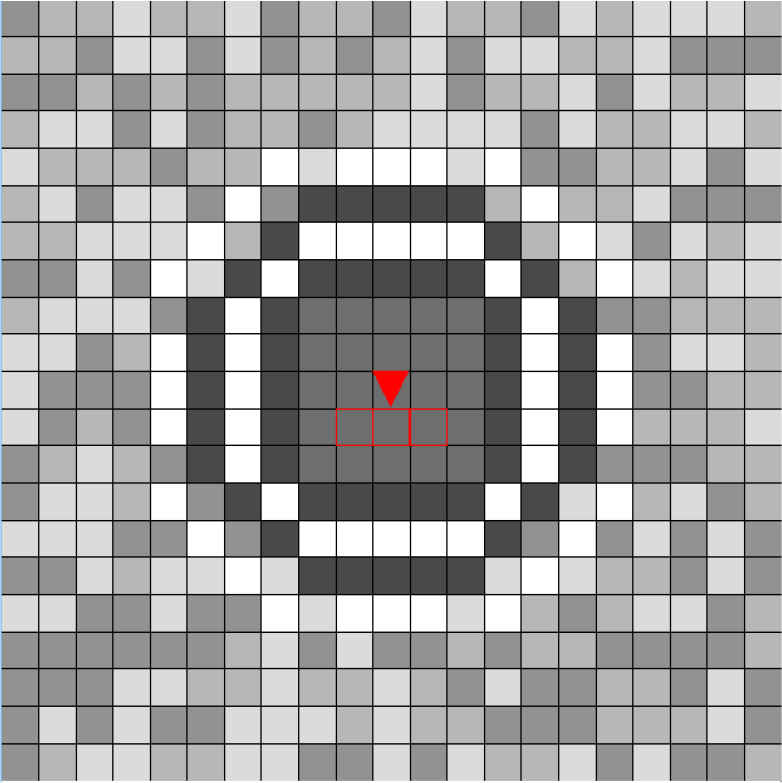
Constructed nest with pufferfish in center.

A nest building video is available at: https://www.youtube.com/watch?v=xOAtLROZma4

The H2Oai machine learning toolkit (https://www.h2o.ai) was used to train a deep learning neural network that successfully learns nest building activity. A typical network has two hidden layers of 200 neurons with a rectified linear activation function. 500 epochs was typically sufficient. In the sample statistics shown in Figure 8, a training dataset of 5 runs under random initial elevation configurations and a test dataset with a different elevation configuration were used. Shown are a graph of training and test errors approaching 0 over time and the test validation confusion matrix.

**Figure 8.**
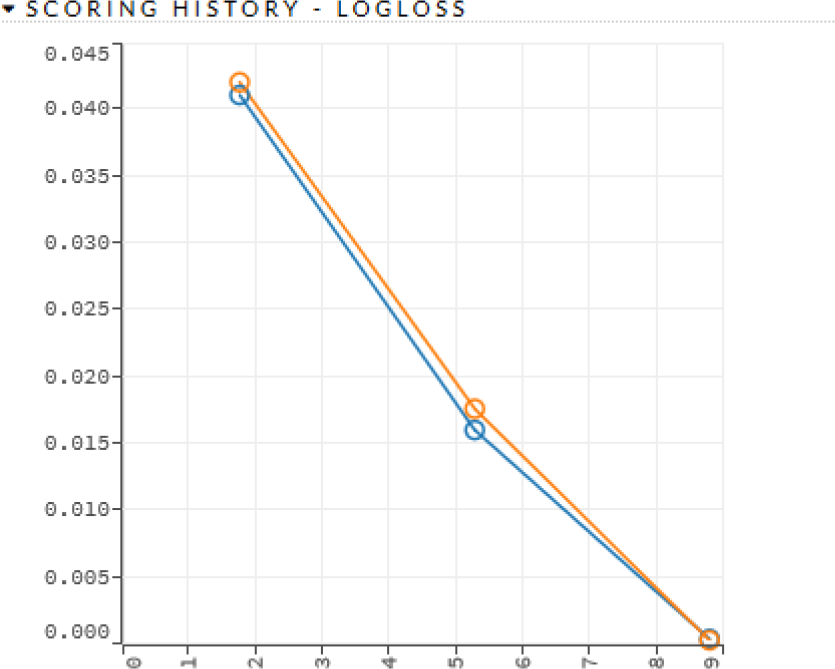
Deep learning neural network statistics.

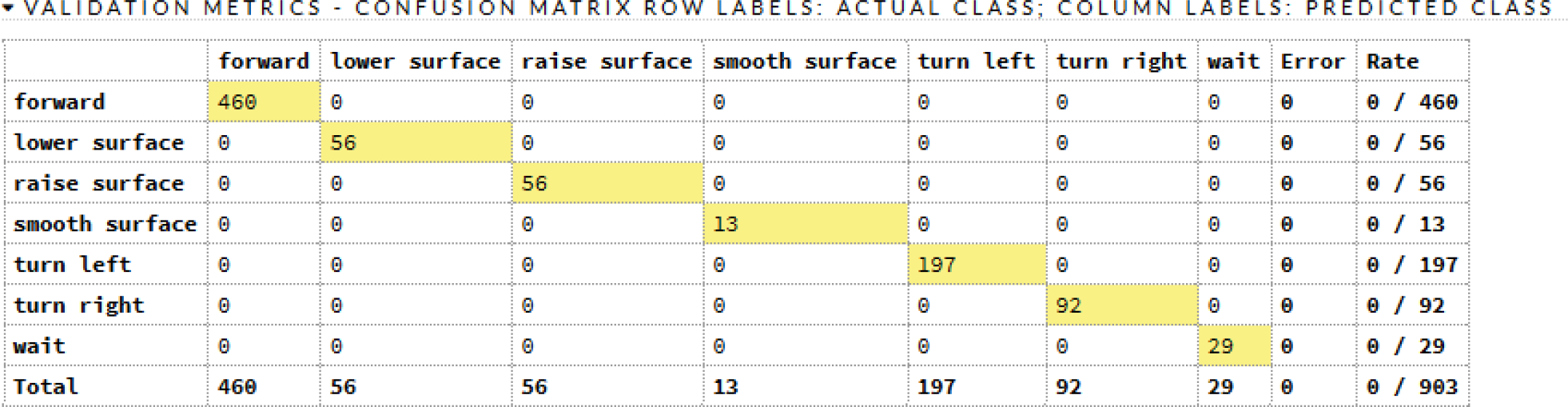

## Conclusion

Building on the replication of C. elegans locomotion and foraging, the pufferfish nest building task represented a further challenge for the Morphognosis model. The result is set of metamorph rules and trained ANN that accomplish the task.

In summary, Morphognosis features:

1. A method for integrating knowledge of events occurring in space and time dimensions in scalable complexity.
2. Nested hierarchies of contexts that provide global-to-local mediating influences.
3. A means of expressing the behavioral interplay of responses and sensory events.

Positive results on the nest building task prompts future investigation. Possible next tasks include:

- Web building. Can a space-time memories of building one or more training webs allow one to be built in a quasi-novel environment?
- Food foraging social signaling. Bees retain memories of foraging food sources that they communicate to other bees through instinctive dancing. Can this task be cast into the model?
- The metamorph structure bears a close resemblance to deep reinforcement learning (Francois-Lavet et al., 2018) training elements, suggesting the possibility of applying such learning to implement goal-seeking behavior.

